# Spatial heterogeneity of cell-matrix adhesive forces predicts human glioblastoma migration

**DOI:** 10.1101/2020.05.06.080804

**Authors:** Rasha Rezk, Bill Zong Jia, Astrid Wendler, Ivan Dimov, Colin Watts, Athina E Markaki, Kristian Franze, Alexandre J Kabla

## Abstract

**Background:** Glioblastoma (GBM) is a highly aggressive incurable brain tumor. The main cause of mortality in GBM patients is the invasive rim of cells migrating away from the main tumor mass and invading healthy parts of the brain. Although motion is driven by forces, our current understanding of the physical factors involved in glioma infiltration remains limited. This study aims to investigate the adhesion properties within and between patients’ tumors on a cellular level and test whether these properties correlate with cell migration.

**Methods:** Nine tissue samples were taken from spatially separated sections during 5-aminolevulinic acid (5-ALA) fluorescence guided surgery. Navigated biopsy samples were collected from strongly fluorescent tumor cores, a weak fluorescent tumor rim, and non-fluorescent tumor margins. A microfluidics device was built to induce controlled shear forces to detach cells from monolayer cultures. Cells were cultured on low modulus polydimethylsiloxane representative of the stiffness of brain tissue. Cell migration and morphology were then obtained using time lapse microscopy.

**Results:** GBM cell populations from different tumor fractions of the same patient exhibited different migratory and adhesive behaviors. These differences were associated with sampling location and amount of 5-ALA fluorescence. Cells derived from weak- and non-fluorescent tumor tissue were smaller, adhered less well, and migrated quicker than cells derived from strongly fluorescent tumor mass.

**Conclusion:** GBM tumors are biomechanically heterogeneous. Selecting multiple populations and broad location sampling are therefore important to consider for drug testing.

**Key points:** GBM tumors are biomechanically heterogeneous

GBM cell migration is inversely correlated with cell-matrix adhesion strength

5-ALA fluorescence intensity during surgery correlates with the motility properties of GBM cells

**Importance of the study:** This is the first study to compare single cell migration and cell-matrix adhesion strength of GBM, using cell lines derived from different tumors and from different regions within the same tumor. Not accounting for internal sampling location within each tumor obscures differences in cell morphology, motility and adhesion properties between patients. Peripherical and marginal tumor cells have different adhesion profiles and are highly migratory compared to those found in the core of the tumor. Aggressive regions of the tumor (highly motile) are linked to the spatial distribution of adhesion strength and are strongly associated with 5-ALA fluorescence intensity. Preclinical tests aimed at developing a treatment for GBM using anti-invasive drugs or adhesion inhibitors, would benefit from using cell lines derived from the tumor periphery (with low 5-ALA intensity) rather than cell lines derived from the tumor core.

## Introduction

Glioblastoma (GBM) is the most common and the most malignant brain tumor. No cure is available despite significant improvements in surgical techniques and radiation technology. The extensive and infiltrative growth pattern of GBM makes surgical resection extremely difficult^1^ and limits the efficacy of radiation therapy by obscuring tumor margins^2^. Tumor recurrence is inevitable; targeted chemotherapy^3^ and immune therapy^4^ have failed to stop tumor recurrence.

Migration and invasion are driven by mechanical forces^5^. The mechanical properties of a GBM tumor and its microenvironment have also shown to contribute to tumor invasion^6–8^. GBM employ a mesenchymal mode of migration^9^ using focal adhesions proteins as molecular clutches to transmit forces to their environments. Targeting integrins or kinases that mediate cell-matrix adhesion has therefore been explored as a strategy to inhibit tumor growth (e.g. focal adhesion kinase (FAK) inhibitor^10^) or to halt the mesenchymal mode of migration employed by infiltrating GBM cells^11^. However, therapeutics designed to target adhesion receptors or proteases have failed in clinical trials^12^, particularly in gliomas (e.g. cilengitide^13^). This failure might be due to the heterogeneity in expression of the adhesion proteins in glioma. Arguably GBM intratumor heterogeneity is the key to understanding treatment failure^14^ and infiltration.

GBM tumors show transcriptomic and genomic distinct subclasses^15, 16^ which vary across patients, but also across individual cells within a tumor^17^. This could explain why gene expression-based molecular classification of brain tumors failed to provide a more accurate prediction of tumor progression and response to treatment. Differential response to treatment is visible during fluorescent guided surgery. GBM cells glow fluorescent pink when 5-aminolevulinic acid (5-ALA) is administered orally prior to surgery. The heterogeneity of 5-ALA-induced PpIX fluorescence observed during surgery was associated with different cellular functions and a distinct mRNA expression profile, where non-fluorescent tumor tissue resembled the neural subtype of GBM and fluorescent tumor tissue did not exhibit a known subtype^18^. However, whether fluorescence heterogeneity is mirrored by physical heterogeneity amongst GBM cells remains unclear.

To investigate the different adhesive and migratory properties of GBM subpopulations, we derived tumor cells from different GBM patients, and from different regions within the same tumor. Tissue samples obtained from the same tumor were collected from spatially distinct locations with different 5-ALA fluorescent intensities. We adapted a microfluidic device for detachment of adherent cells through shear stress^19, 20^. We demonstrate substantial intra- and inter-tumoral heterogeneity; adhesion strength varied tenfold (from 15 Pa to 150 Pa). Cells from the weak fluorescent tumor rim and non-fluorescent margins were smaller, less adherent with highly migratory behavior, suggesting that differential adhesion and migration speed between subpopulations of cancer cells may contribute to tumor invasion. Such heterogeneity could also explain recently observed differential responses of patients to adhesion-blocking drugs.

## Materials and Methods

### Sample collection

Three patients newly diagnosed with GBM underwent surgical resection. Aminolevulinic acid (5-ALA) fluorescence was orally administered four hours before induction of anesthesia at a dosage of 20 mg/kg. Three different regions within the tumor were biopsied. Six tissue samples from two different patients were taken from spatially separated sections using MRI stealth imaging. Navigated biopsy samples were collected from strongly fluorescent tumor cores, a weak fluorescent tumor rim, and non-fluorescent tumor margins (Figure 1A). In addition, three tissue samples from a third patient were collected from similar locations (results described in supplementary material). Tissue collection protocols complied with the UK Human Tissue Act 2004 (HTA license ref. 12315) and have been approved by the local regional ethics committee (LREC ref. 04/Q0108/60).

**Figure 1:**
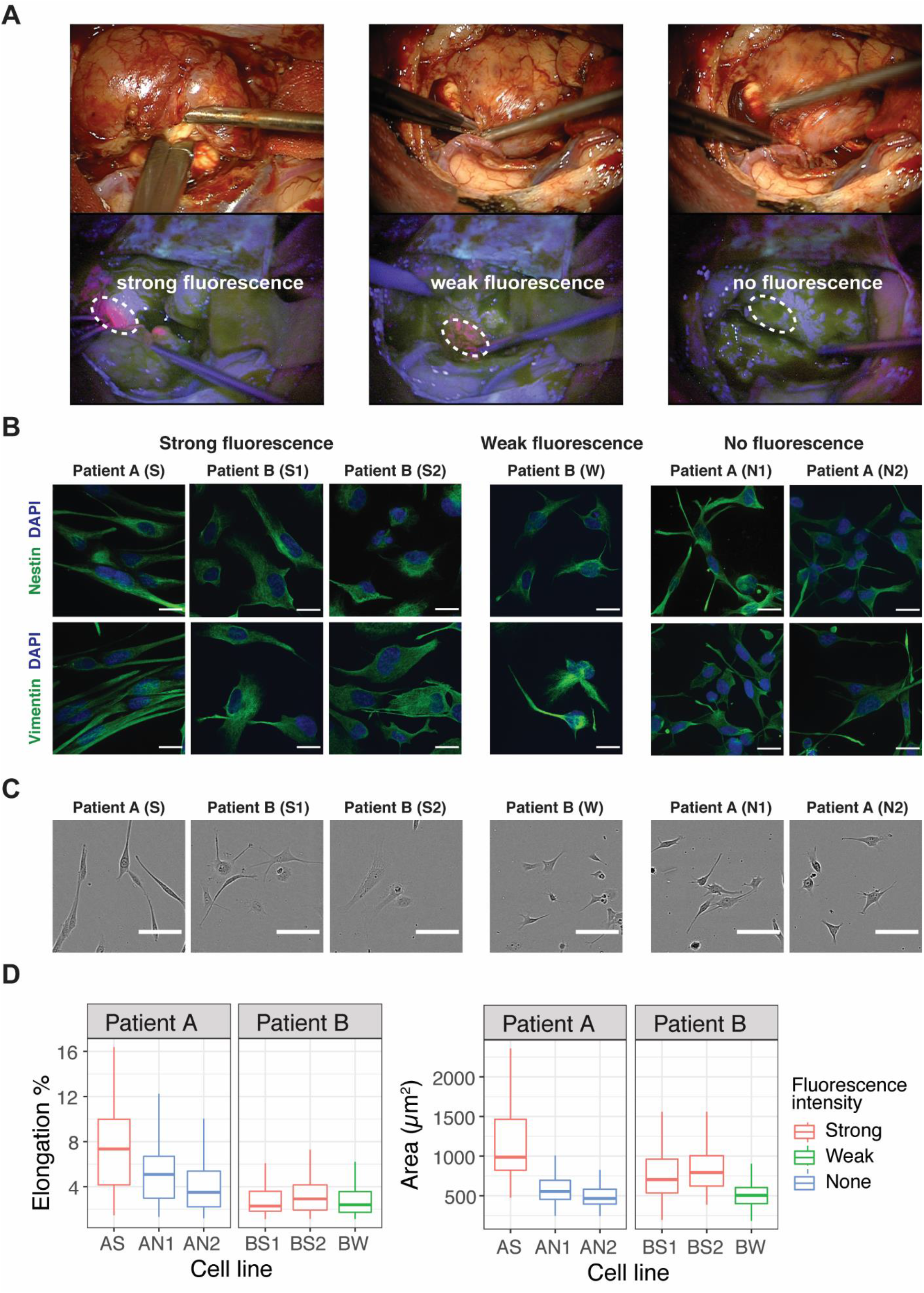
Fluorescence and Morphological Characteristics of GBM Patients Derived Cell Lines. (A) White light image of resection cavity and corresponding fluorescence view of surgical field demonstrating the heterogeneous fluorescence pattern of 5-ALA (bright strong pink from tumor core, faint/weak pink from tumor rim, and non-fluorescent tumor margin). Intraoperative images were obtained via ZEISS OPMI PENTERO 800, with 39x magnification. We collected three strongly fluorescent tissue samples from the tumor mass (S from patient A, S1 and S2 from patient B), two non-fluorescent tumor margins (N1 and N2) from patient A, and one weak-fluorescent tissue sample (W) from patient B. (B) Immunofluorescence staining for evidence of GBM stem like cells. Neural stem cell marker nestin (green), and vimentin (green) are expressed in each cell line; scale bars 20 μm. (C) Representative phase-contrast images showing cells derived from strong fluorescent tissue samples has larger area compared to cells derived from weak- and non-fluorescent samples. Patient A exhibited a more elongated morphology compared to patient B (***p<0,.001); scale bars 100 μm. (D) Morphological quantification of cells cultured on PDMS substrates of ~1.5kPa stiffness (n > 100 cells). Boxplots show spatial and intertumoral heterogeneity in cell area and cell elongation. Boxplots represent the “minimum”, first quartile (Q1), median, third quartile (Q3), and maximum.

### Derivation of GBM stem-like cells

Cell derivation and maintenance have previously been described^21^. Briefly, tissue was mechanically minced and enzymatically dissociated before passing through a 40 μm cell strainer. Cells were seeded in serum-free medium (SFM) (phenol red free Neurobasal A) with 2mM l-glutamine and 1% volume/volume (v/v) penicillin/streptomycin (PS) solution with 20 ng/mL human epidermal growth factor, 20 ng/mL zebrafish fibroblast growth factor (FGF-2), 1% v/v B27 SF supplement, and 1% N2 SF supplement. Cells were allowed to form primary aggregates. Spheroid aggregates were collected and plated, onto Engelbreth-Holm-Swarm sarcoma extracellular matrix (ECM, Sigma)–coated flasks (ECM 1:10 dilution with HBSS) and allowed to form a primary monolayer. When the primary monolayer reached 80% confluency, cells were passaged to generate the subsequent monolayers by mechanically and enzymatically dissociating remaining aggregates. Cells were maintained at 37°C and 5% CO2.

### Immunocytochemistry

Cells were seeded in duplicates into μ-Dish 35 mm (60,000 cells/ibidi dish). After 48 hours, cells were washed with sterile phosphate buffered saline (PBS, Thermo Fisher) before being fixed in 4% paraformaldehyde for 30 minutes (10 minutes at 4°C followed by 20 minutes at room temperature). Cells were washed and then permeabilized for 4 minutes in 0.2% Triton in PBS. Cells were washed again and incubated in blocking buffer (1% bovine serum albumin (BSA), and 2% normal goat serum (NGS), in PBS) for 1 hour at room temperature before incubating with the primary antibody for 2 hours. The following primary antibodies were used: Nestin (Abcam, ab22035, 1:100), Vimentin (Abcam, ab8069, 1:200), NG2 (Abcam, ab83178, 1:200), Ki67 (Abcam, ab16667, 1:200), Vinculin (FAK100, Sigma 1:200), F-actin (FAK100, Sigma 1:500). Secondary antibodies were applied for 1 hour (goat anti-mouse Alexa Fluor 488 pre-adsorbed abcam, 1:750 dilution and goat anti-rabbit Alex Fluor 594 pre-adsorbed abcam, 1:750). Nuclei were stained with DAPI (Roche)

### Immunoblotting

Cell lysis was performed using cOmpleteTM, EDTA-free lysis-M buffer with protease inhibitor (Roche). Protein concentrations were determined using PierceTM BCA kit (Thermo ScientififTM). Equivalent amounts of protein were electrophoresed on SDS-polyacrylamide gels. The gels were then electroblotted onto PVDF membranes. After blocking with Odyssey Blocking Buffer (TBS) (LI-COR), membranes where incubated with the primary antibody overnight (Paxilin, Vinculin, FAK, GAPDH (codes need to be looked up at home). Finally, the relevant protein was visualized by staining with the appropriate IgG H&L secondary antibody coupled to either IRDye^®^ 800CW or IRDye^®^ 680RD. The antigen of interest was detected using the LI-COR Odyssey CLx Infrared Imaging System. Results were analysed using ImageStudio.

### Poly-di-methyl-siloxane (PDMS) substrates for studying cellular motility and morphology

NuSil GEL-8100 (NuSil) was prepared in a 1:1 ratio of component A and component B and mixed well for 60 seconds. 1% (w/w) 10:1 (base/crosslinker w/w) Sylgard-184 (VWR) was added to the GEL-8100 and mixed well for 60 seconds. For cell morphology experiments approximately 120mg of the PDMS was added to μ-Dish 35 mm (120mg/dish). For cell migration experiments 80mg/well of PDMS was added to 24-well culture plates (Corning Life Science, Tewksbury, MA, USA). Coated vessels were baked at 65°C for 13 hours. This treatment gave a shear modulus value of G =1.53±0.12 kPa, n = 16, verified by atomic force microscopy (AFM) indentation. The AFM setup consisted of a JPK CellHesion 200 scanner, and the indentation probe was made by gluing a spherical polystyrene particle (90 um diameter), to a tipless AFM probe (SHOCON-TL, k ~ 0.1 N/m). Vessels were sterilized by immersion in 70% (v/v) ethanol in distilled water for 15 minutes, followed by two rinses with PBS. The PDMS surface was coated with ECM at a concentration corresponding to a surface density of 6.67 μg/ml assuming complete adsorption.

### Cell detachment assay

Microfluidic devices were manufactured (Supplementary information S1) and sterilized by immersing in 70% ethanol in distilled water (v/v) in a Petri dish, and perfusing with the same liquid at 10 mL/h for 30 minutes. The devices were lifted from the ethanol and perfused with phosphate-buffered saline (PBS, Thermo-Fisher) at 5 mL/h for 1 hour to rinse the ethanol and allow PBS to permeate the bulk of the device. The devices were filled with 200 μg/mL ECM matrix and placed in the incubator overnight (approximately 16 hours). SFM was equilibrated overnight in the incubator to minimize bubble formation. Cells were detached as described previously and resuspended to a concentration of 3.5×10^^6^ cells/mL. Cells were loaded into 1mL syringes and perfused into the devices at 30 μL/h for 5 minutes. Perfusion was resumed for 3 minutes with the inlet and outlet reversed to seed cells evenly. The devices were placed in the incubator for 4 hours to promote cell attachment. The devices were then perfused for 20 hours with equilibrated SFM at a flow rate of 10 μL/h to allow cells to fully spread^22–24^. For cell detachment, neurobasal Medium without supplements was also equilibrated overnight in the incubator to minimize bubble formation and to control dissolved gas concentration and pH. Cells were subjected to a steady flow rate of 0.16 mL/min, 0.33 mL/min and 0.5 mL/min to create a constant shear force on the cell (Supplementary information S1).Phase-contrast images of a single field of view were taken every two seconds at 160×magnification (Zeiss Axio Observer.Z1). The maximum flow rate used was 0.5 mL/min, (a shear stress of 75.96 Pa), although shear stress of up to 506 Pa is achievable with the system, limited by the maximum flow rate of the syringe pump.

### Time lapse measurements and cell tracking analysis

Cells were cultured in 24-well culture plates (Corning Life Science, Tewksbury, MA, USA) according to the manufacturer’s instructions and visualized using a real-time cell imaging system (IncuCyte™ live-cell ESSEN BioScience Inc, Ann Arbor, Michigan, USA). Ten thousand cells/well were cultured on PDMS substrate of 1.5 kPa stiffness. Cells were seeded in triplicates and imaged every 10 min for 48 hours. Time-lapse images were acquired with IncuCyte, a live cell imaging microscope.

Raw images were processed in CellProfiler^25^ to detect cell outlines. Cells were tracked using automated tracking in TrackMate^26^. Cell area was calculated after 24 hours, using the hierarchical K-means thresholding module in Icy^27^. The mean square displacement (MSD) of each cell from its starting position was calculated. The MSD of actively moving cells should be larger than the expected MSD of a freely diffusing (Brownian) particle of comparable size:

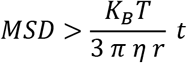

where *K_B_* is the Boltzmann constant, *T* = 310 K the absolute temperature, *η* the viscosity of the medium, *r* the radius of the cell and t the time interval. We assumed the viscosity of the medium to be *η* ≈ 0.78 mPa s.

### Statistical analysis

Data were verified from at least two independent biological experiments. For experiments involving single-cell analysis, n ≥ 100 cells. The order of data collection was randomized; no blinding was performed and no data were excluded from the analysis.

Morphological differences between patients and fluorescence groups were tested using linear regression. First, we tested differences between patients and differences between fluorescence groups independently. Then we tested for differences simultaneously using an additive model (for patients and fluorescence intensity). Since there was no statistically significant difference between weak- and non-fluorescent cell lines but both were different from strongly fluorescent cell lines, we pooled the weak- and non-fluorescent groups. To check whether sampling from multiple locations violated our standard linear model assumptions, we tested unexplained heterogeneity using mixed effect regression. Including a random effect of all lines did not significantly improve model fit. This indicates that a mixed effect model is not required.

Adhesion heterogeneity between cell lines was analyzed using Welch’s *t*-test (unequal variance) with Bonferroni correction (15 comparisons). The shear stress required for 50% cell detachment was found by performing an orthogonal distance regression, and the resulting means and standard errors were used in the Welch’s *t*-test formula. Differences in cell migration between patients and fluorescence groups were tested using weighted linear regression (using MSD slope estimate as the outcome). First, we tested differences between patients and differences between fluorescence groups independently. After pooling the weak- and non-fluorescent cell lines, we tested for differences simultaneously using an additive model (for patients and fluorescence intensity).

Statistical analyses and plotting were performed in Python 3.6, MATLAB or R statistical software package.

## Results

### Relationship between tissue fluorescence heterogeneity and GBM cell morphology

Primary GBM cell lines maintained in serum-free media expressed neural stem cell markers nestin and vimentin (Figure 1B). This confirmed the presence of glioma stem-like cells which are regarded as tumorigenic and associated with the heterogeneity in GBM^28^. We then investigated the cell spreading area which is known to be an indicator of how cells mechanically interact with their extracellular environment^29^. Cells were cultured on polydimethylsiloxane, with a low modulus consistent with the stiffness of the environment that GBM infiltrate, which ranges from 0.1 kPa to 10 kPa^30^. Glioma cells are known to spread on substrates of stiffness around 1kPa^31^However, on extremely compliant substrate (~150 Pa) GBM cells exhibit rounded morphologies with diffuse distributions of F-actin that are unable to migrate productively^32^. To compare between lines, we used a single stiffness, ~1.5kPa (Methods) (Figure 1C).

To explain the variability in cell morphology observed among patients’ cell lines (Figure 1D), we first tested whether cells from patient A differed from those of patient B. Cells derived from patient A exhibited a more elongated morphology compared to cells derived from patient B (***p < 0.001), but there was no significant difference in cell area. However, cells derived from strongly fluorescent lines differed significantly in cell area and cell elongation between patients (***p < 0.001). We therefore adjusted our model to account for fluorescence intensity when comparing between patients (Methods). We found that cells from patient A were both larger and more elongated than cells derived from patient B (***p<0.001).

The results demonstrate morphological differences within each tumor and between tumors and show that this heterogeneity is related to 5-ALA fluorescence intensity. Not accounting for this variability can obscure the difference between patients. To investigate whether the differences in cell morphology relate to the way cells adhere to their environment, we explored whether heterogeneity in cell area and shape correlate with specific patterns of key cytoskeletal proteins.

### The organisation of actin filaments and vinculin suggests intertumoral heterogeneity in GBM cell-matrix adhesion

Actin filament disassembly is required for cell spreading, and the binding to proteins such as paxillin and vinculin is necessary for focal adhesion development^33^. We imaged the localization and structure of actin filaments and the actin binding protein vinculin. The shape and distribution of actin and vinculin differed between cells derived from strongly fluorescent lines and cells derived from weak- and non-fluorescent lines. The colocalization of vinculin with F-actin was evident at the edge of cells derived from strongly fluorescent lines (Figure 2A). Adhesion proteins paxillin, vinculin and Focal Adhesion Kinase (FAK) were expressed across all cell lines (Supplementary Figure S2), but their expression levels did not reflect the arrangements and assemblies revealed in immunocytochemical images. The binding of vinculin to F-actin suggests spatial differences in focal adhesion assembly and enlargement within the tumor. We therefore built a set-up to quantify the intra-and-inter tumoral heterogeneity in cell-matrix adhesion.

**Figure 2.**
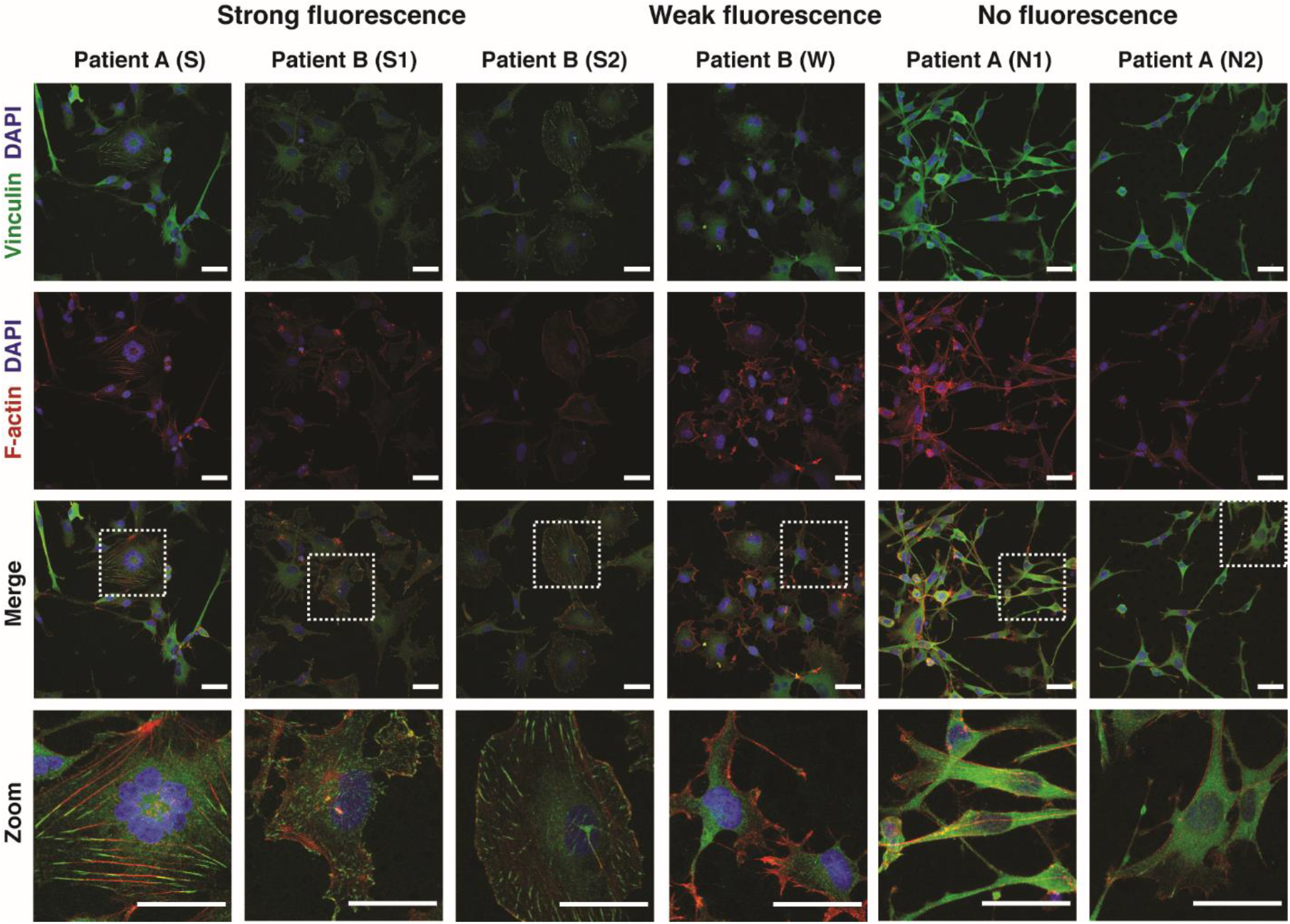
F-actin cytoskeleton and Vinculin proteins expression in GBM lines. Confocal images showing the colocalization of vinculin and F-actin at the edge of cells derived from strong fluorescent lines compared to cells derived from weak- and non-fluorescent lines. Cells were stained with TRITC-phalloidin (red) to visualize the actin cytoskeleton, vinculin (green) and DAPI (blue). Scale bars 40 μm.

### Cell-matrix adhesion heterogeneity within and between patient-specific tumor samples

To measure adhesion strength, we built a microfluidic device that supported overnight culture of GBM cells and generated sufficient shear force to trigger their detachment (Supplementary Figure S1). Patients’ derived cells were delivered into the channels via syringe pumps (3.5×10^6^ cells /mL). Cells were gently perfused with serum free medium and placed in an incubator for 24 hours to allow adhesion to fully establish and cells to fully spread^22–24^. Cells were subjected to a controlled flow rate to create a constant shear force on the cell (Supplementary information S1). The fraction of detached cells was measured over time (Figure 3A). We consistently found an initial phase of rapid detachment followed by a second phase of slower detachment (Figure 3B), consistent with previous parallel-plate and microfluidic detachment assays^34, 35^. This transition manifests itself in most experiments as a “knee” in the time-detachment curve after the first 5 to 10 minutes of exposure to the shear force (Supplementary Figure S3).

**Figure 3:**
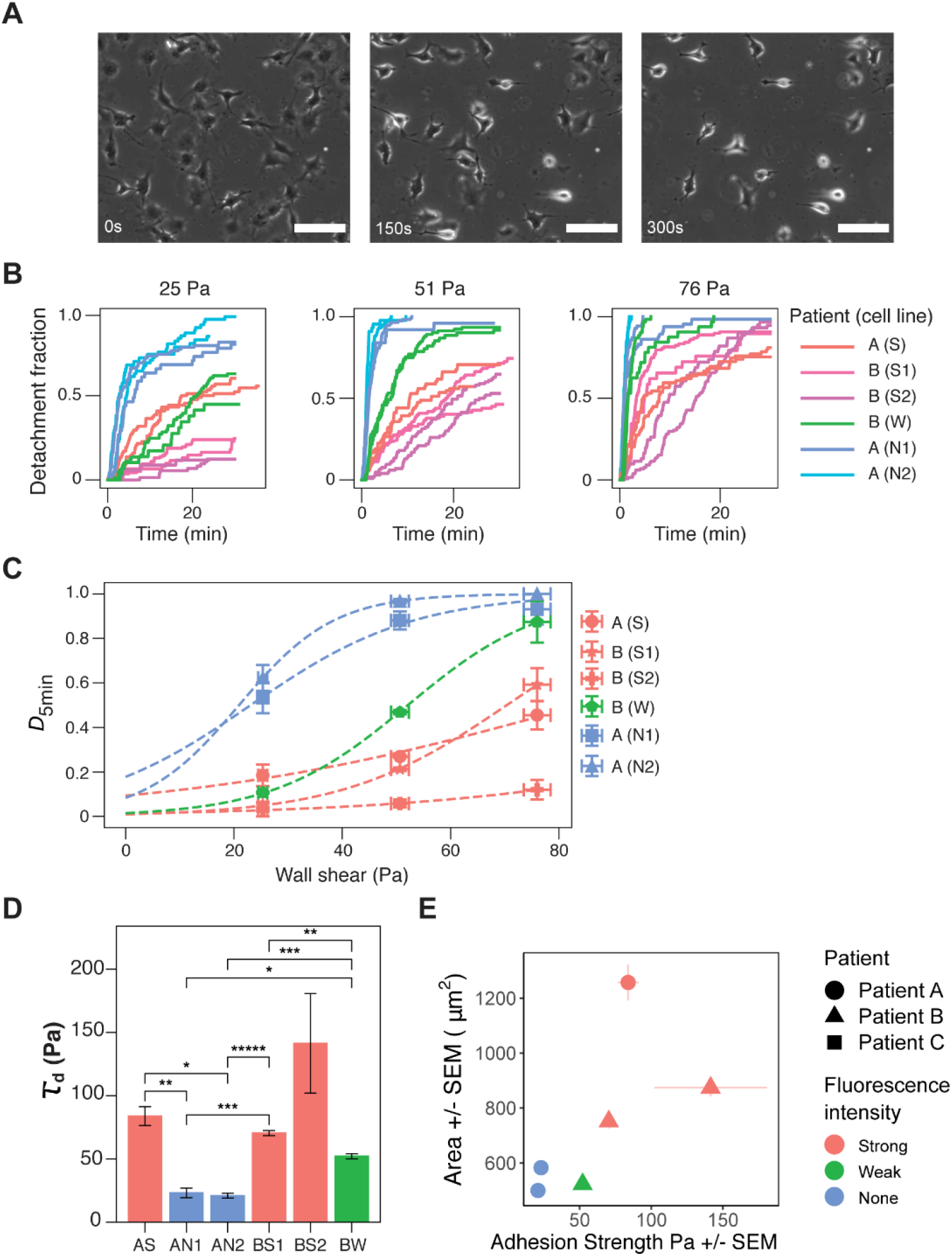
GBM exhibits spatial and inter-tumor heterogeneity in cell-matrix adhesion. (A) Representative phase-contrast images of cells under microfluidic shear detachment at 0, 150, and 300 seconds. Here BW cells are exposed to 0.33 mL/min of flow, corresponding to 51 Pa of shear stress. Scale bars 100 μm. (B) Time-detachment profiles for each of six lines surveyed (n=2, 30-140 cells per experiment) for a range of shear stresses. Detachment increases with shear stress, but the relative ordering of the lines remains constant except for cell line AS. Detachment initially occurs rapidly, and plateaus for most cell lines. (C) The inflection point of each sigmoidal curve was extracted to define τ_*d*_, a measurement for the cell-matrix adhesion strength of the cell lines. (D) Differences between weak- and strongly-fluorescent lines were statistically significant for all intra-patient pairs except BW and BS2 (one * indicates (*p <0.05) and additional *s indicates further orders of magnitude, under Welch’s t-test with Bonferroni correction). The mean value of τ_*d*_ for patient B was significantly higher than that of patient B (**p = 0.005). (E) Relationship between cell morphology and cell-matrix adhesion strength. Strong fluorescent cells are more adherent and larger than weak- and non-fluorescent cells. No clear relationship between adhesion strength and cell elongation.

There is some evidence suggesting that the second phase is a result of adhesion weakening in response to imposed flow, driven by Rho regulation of focal contact maturation and turnover^36, 37^. As a consequence, cell detachment measurements taken later in time may not be representative of cell-matrix adhesion under normal condition. We chose to take measurements of detachment at 5 minutes of flow, as a quantification of adhesion strength (Supplementary Figure S3).

Detachment at 5 minutes was expressed as a fraction of the cell number at t=0. Detachment fraction as a function shear stress was fitted to a logistic function, and the shear stress required for 50% detachment (τ_*d*_) was identified. as the inflection point of the logistic fit (Figure 3C). τ_*d*_ ranged between 15 Pa and 141 Pa. This range is consistent with microfluidic single-cell adhesion strength measurements of 3T3 fibroblasts, which fell between 20 and 220 Pa^20^.

Cells derived from weak- and non-fluorescent cell lines had significantly lower cell-matrix adhesion than cells derived from strongly fluorescent cell lines across patients A and B (*p < 0.05) (Figure 3D). The mean value of detachment strength of patient B was significantly higher than that of patient A (**p = 0.005). The data shows that both intra- and inter-tumoral cell-matrix adhesion heterogeneity is present within GBM. Intratumoral cell proliferation is strongly associated with the fluorescence intensity of samples, and hence the spatial density and distribution of cells. Strongly fluorescent cells derived from tumor core were more adherent to the extracellular matrix, and had a larger cell spreading area (Figure 3E). Weak- and non-fluorescent cells had smaller spreading area and were less adherent.

### Migratory behavior of tumor derived cell populations is predicted by adhesive forces

To quantify the migratory behavior of GBM cells, we recorded their trajectory on a compliant PDMS substrate. Their movement is stochastic at the scale of 10 mins and is best characterized by an effective diffusion coefficient D. which capture the rate of change of the squared end-to-end distance travelled (Mean square displacement (MSD) Δ*R*^2^) as a function of time interval Δ*T: ΔR*^2^ = D Δ*T*. The larger D, the more migratory the cell line is.

We first tested whether cells from patient A differed in motility from patient B and found no significant differences in their diffusion coefficient. After adjusting our model to account for fluorescence intensity (see Methods), we found that cells from patient B migrated faster than patient A (*p<0.05). Cells derived from strongly fluorescent tissue samples were significantly slower than cells derived from weak- and non-fluorescent cell lines (**p < 0.01) (Figures 4A).

**Figure 4.**
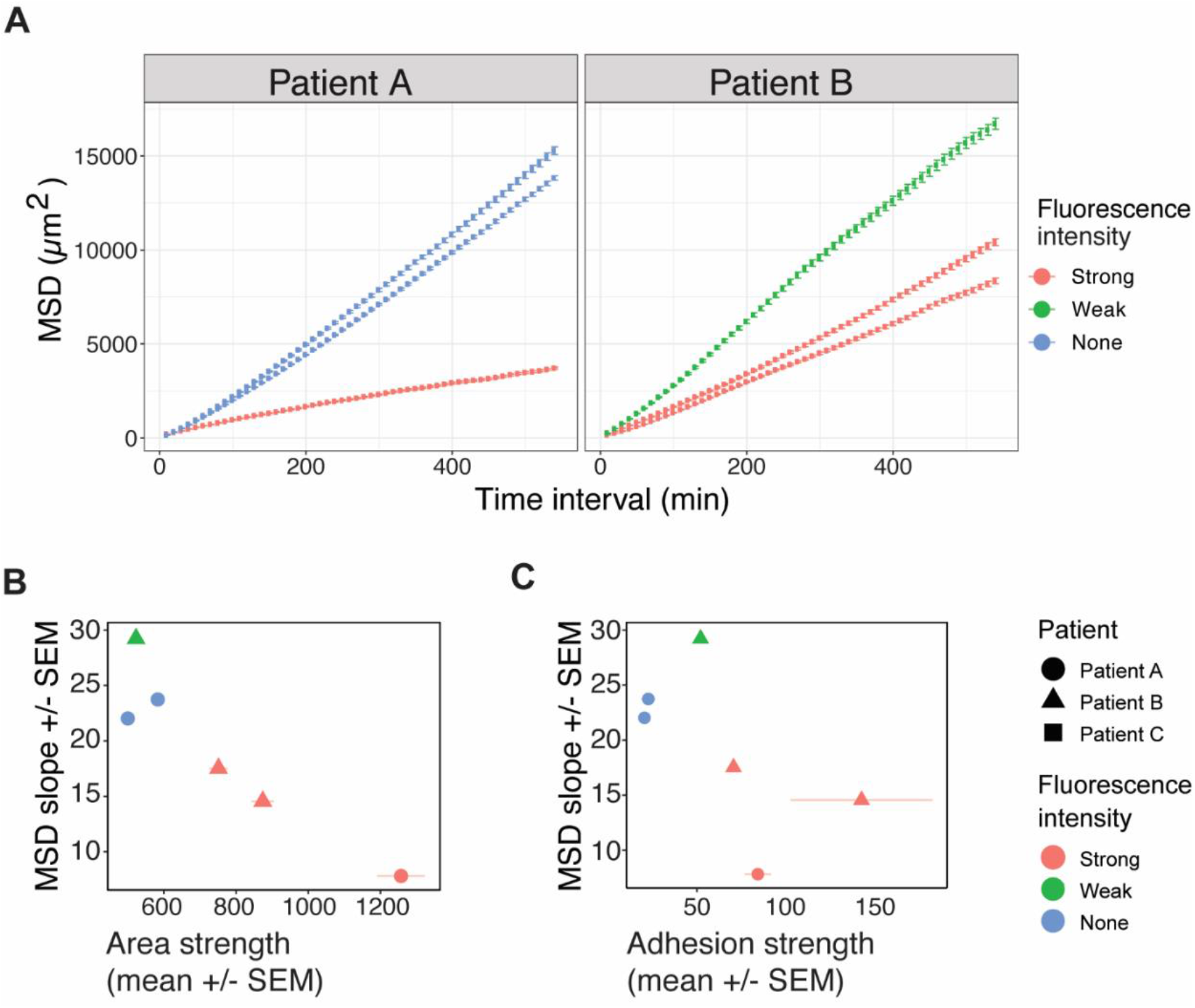
Cell-matrix adhesion is an important factor for GBM cell migration. (A) MSD as a function of time interval (Δt) calculated from cell trajectories with vertical bars representing standard errors (more than 100 trajectories per cell line from two biological replica). Having accounted for the cell line fluorescence intensity, we found cells from patient B migrated faster than patient A (*p<0.05). Cells derived from strongly fluorescent tissue samples were significantly slower than cells derived from weak- and non-fluorescent cell lines (**p < 0.01). (B) Cells with larger cell area migrate less than those with smaller cell size derived from the weak- and non-fluorescent samples. (C) Cells derived from strong fluorescent tissue samples are more adhesive and less migratory.

Cells derived from strongly fluorescent core lines had a larger area, migrated less (Figure 4B) and adhered more than cells derived from weak- and non-fluorescent cell lines (Figure 4C). This result is consistent with previous models of pseudopodial migration, in which a certain level of adhesion is required for generation of traction forces, but excessive adhesion impedes motion^38^. Additionally, the less adherent and more migratory GBM cells are also from weak- and non-fluorescent cell lines, derived from tumor rim and margin respectively. This observation suggests a potential mechanism of differential diffusion. Both cell migration and cell-matrix adhesion strength were associated with cell size (Figure 3B and Figure 4B) and not cell shape (Supplementary Figure S4).

Our results also show significant differences between patients and that cell-adhesion strength could be accurate predictor of tumor cells migration unlike molecular classifications which as yet have failed to provide accurate prediction.

## Discussion

In order to invade the surrounding tissue, GBM cells have to exert forces on their environment and mechanically interact with it. They also have to undergo biological and morphological changes that allow them to migrate through the perivascular spaces and white matter tracts of the brain^11^. Instead of observing the tumor tissue *in* vivo, our approach isolated cell lines and measured their adhesive and migratory properties under controlled conditions allowing direct comparison. The results are therefore not confounded by any spatial differences in extracellular matrix remodelling, and in nutrient or signalling factor concentration. Our findings establish that the variation of GBM cell-matrix adhesion correlate well with GBM cell area and cell migration.

Cells derived from tumour rim and tumor margins are smaller, adhere less well, and migrate quicker than cells derived from core tumor mass. Characterising cell–ECM interactions alongside cell motility provides a better picture of the biomechanical context^39^. The relationship between cell-matrix adhesion, morphology and migration was further confirmed by analysing a third patient whose tissue samples were obtained from three similar locations in the tumour; the derived cell lines exhibited similar morphology (Supplementary Figure S5), adhesive (Supplementary Figure S6) and migratory behaviour (Supplementary Figure S7). Cells size was directly correlated with cell-matrix adhesion and inversely correlated with cell motility. This also highlights that it is important to account for intratumor heterogeneity before comparing between patients’ lines.

The fact that adhesion strength is associated with reduced movement^40^ in vitro suggests a simple model where migratory forces must overcome adhesion forces to generate motion. This idea was tested numerically (Supplementary information S2) in order to relate MSD curves with the measured values of cell adhesion. This approach can be extended to simulate mixed populations of core and marginal cells. The variation in adhesion strength leads to variations in amount of migration, causing cells with lower adhesion to be predominant at the leading edge, consistent with the clinical observation (Supplementary Figure S8). Differential migration is known to contribute to cell segregation^41^ and may play a role in the spatial distribution of cells in tumor.

The intratumor heterogeneity was associated with sampling location and the heterogeneity of the 5-ALA induced fluorescence observed during fluorescence-guided resections. Recently it has also been shown that fluorescent and non-fluorescent tissue samples can be distinguished genetically^18^. Due to our controlled experimental conditions, much of the observed intra- and inter-tumoral heterogeneity is likely to of genetic or epigenetic origin, rather than caused by spatial differences in the original tissue’s microenvironment. Our results are consistent with a broader picture of GBM as a genetically heterogeneous cancer^14, 17^, and genetic changes in genes crucial in cell mechanics may be implicated in tumor progression.

This intra-inter tumor heterogeneity could explain recently observed differential responses of patients to adhesion-blocking drugs, and strongly suggests that different parts of the same tumors should be treated. The failure of adhesion-block treatments may be due to the observed intra-and intertumoral heterogeneity, as typically preclinical tests use only one or two cell lines, which may not be representative of the distribution of integrin expression and adhesion in GBM tumors. Given that integrins play a crucial role in brain tumor infiltration^42^ and GBM intratumor variability in integrin expressions^43^. New trials that aim to halt GBM recurrence by inhibiting radiation-induced invasion gains and signaling changes^11, 44^, or kinase inhibitors^10^, could benefit from accounting for intra-and-intertumoral differences in single cell migration.

## Notes

**Funding**: This research was supported by Engineering Clinical Practice Grant and the European Research Council (Grant No. 240446 and Consolidator Grant No. 772426). Financial support for RR was provided by Nabaa El Mahabaa, Egypt, Gonville & Caius College and the European Research Council.

**Conflict of interest**: The authors report no conflict of interest, and the present study has not been presented elsewhere.

### Competing Interest Statement

The authors have declared no competing interest.

