## supplementary_methods_table_figures for "Spatial heterogeneity of cell-matrix adhesive forces predicts human glioblastoma migration"

### Supplementary Information (S1)

#### Microfluidic device manufacture

Molds were made using standard methods for photolithography in SU-8 2025 (Micro-chem). Photomasks were inkjet printed in black on clear transparencies (HP). In darkness, SU-8 was spin-coated onto a polished glass slide (Corning Polished Glass 7059, Apex Optical Services) for 30 seconds at 500 revolutions per minute (RPM) and 60 seconds at 2000 RPM (VTC MTI-100). The sample was soft-baked on a hot-plate at 70◦C for 5 minutes, and 100◦C for 15 minutes. The sample was air-cooled and the photomask was placed printed-side down onto the baked SU-8. The sample was cross-linked in an ultraviolet light exposure chamber for 60 seconds. The sample was baked again at 70◦C for 60 seconds and 100◦C for 5 minutes. The sample was air-cooled and placed in Microposit EC solvent (Dow Chemical) for 5 minutes. The finished mold was rinsed using isopropanol and stored in a Petri dish. Sylgard-184 (VWR) was prepared in a 10:1 base/curing agent (w/w) ratio and de-gassed. The mixture was poured over the mold and baked at 65◦C for 4 hours. The PDMS stamp was peeled from the mold and devices were cut out using a scalpel. Inlet and outlet ports were made using a 1mm biopsy punch. Glass slides were cleaned with isopropanol and compressed air. The devices and glass slides were placed in a plasma chamber (Harrick) on high setting for 60 seconds, gently pressed together, and baked at 60◦C for 2 hours. The ports of finished devices were protected with tape until use.

#### Determination of Shear Stress Exerted on Cells in Response to Flow

Assuming a laminar, incompressible, and linear viscous flow through the channel, a differential equation governing the relationship between pressure drop and shear stress can be derived using a force balance on fluid elements. The equation can in turn be integrated over space to determine the relationship between these quantities and the flow rate through the channel. Various approximations for a rectangular channels are possible[1-3], but the simplest assumes that the channel is adequately represented by parallel plates, ignoring the side walls. This assumption is valid because the width of the channel is large relative to the height (20x). The resultant 1D problem can be easily solved to give an expression that determines the shear stress over the height of the channel as a function of flow rate:

$$\frac{\Delta P}{L} = -\frac{12Q\mu}{h^3 w}$$

$$\tau = \frac{h\Delta P}{2L}$$

where $\Delta P$ is the pressure drop (Pa) over the length of the channel $L$ (m), $Q$ is the volumetric flow rate ($m^3/s$), $\mu$is the dynamic viscosity (Pa s), $h$ is the channel cross-sectional height (m), $w$ is the cross sectional width (m), and $\tau$ is the shear stress at the bottom and top channel walls (Pa). At the flow rates exerted, the Reynolds number was checked to ensure that the laminar flow assumption held (Re < 2000). We additionally assumed that the shear stress at the bottom wall of an empty channel was the same as that exerted on the interface between cell and matrix. The shape of cells may alter the effective cross-sectional area of the channel, and may induce an additional body drag due to pressure differences across the length of the cell. However, previous works have demonstrated that these factors have negligible effect on the forces exerted on the cell in the relevant flow regimes[2] .

Time lapse images were processed using MATLAB to determine detachment events in response to flow. Briefly, a local entropy mask was applied to every frame and thresholded to segment image regions as cell candidates. Masks adjacent in time were subtracted to identify candidate detachment events, and these were manually validated to generate time-detachment curves for each experiment.

To identify the shear stress required to detach a particular cell line, we considered the time-detachment plots from each time-lapse video. The detachment typically increased rapidly at first and slowed down, so we sought to identify the "knee" that marked this transition using two independent methods. First, we took linear fits of the data using increasing time windows. Assuming the effect of the transition between the two phases of detachment is larger than any higher-order relationships between time and detachment in the first phase, the goodness of fit should decrease as the "knee" is included. We measured the $R^2$ value of the best linear fit as a function of the time window for each experiment, and the maximum for the average was found to be at 8.74 minutes (S2A). In the second method, we identified the "knee" using the Kneedle algorithm, based on characterization of the first and second derivatives of detachment percentage over time[4] . According to this algorithm, only 25% of experiments had a "knee" before 7.5 minutes (Supplementary Figure 3B). We selected a timepoint of 5 minutes for all cell lines to be well-within the rapid detachment regime.

Detachment at 5 minutes was expressed as a fraction of the cell number at t=0. Detachment fraction as a function of flow rate and hence shear stress was fitted to a logistic function, and the shear stress required for 50% detachment ($\tau_d$) was identified as the inflection point of the logistic fit with minimum mean squared error (Figure 3C). This value of shear stress was used as a comparative metric of detachment stress between cell lines. To verify the suitability of this approach for range of flow rates required to examine GBM adhesion, we tested the detachment of cell line BS2 at its predicted =141.7 Pa (*n* =2), and observed a detachment percentage of 58.5% ± 22% (± STD).

**Cell Detachment Detection for Microfluidic Assay**

A cell counting algorithm called Semi-automated Local Entropy Difference (SLED) was developed to register individual detachment events from a time series of phase-contrast images taken every two seconds.

First, a suitable analysis area is extracted from raw images. A binary mask containing the walls of the channel is obtained by Otsu thresholding based on brightness and any thresholded cells were excluded based on an assumption that only the relevant features (i.e. the walls) touched the edges of the image. The mask is then downsampled and fitted to the equations of a set of parallel lines using linear least squares. The slope of these lines is used to identify any orientation differences between images due to channel manufacture or movement of the microscope stage, and the images were rotated appropriately. The intercepts are then used to identify the position of each of the channel walls. The middle 80% of the image is taken to prevent counting of cells experiencing wall effects and to eliminate artifacts in image processing produced by the bright channel walls.

The initial number of cells is manually counted by the user on the first frame of the image sequence. The images are then filtered based on the local entropy of a 7x7 neighborhood centered at each image pixel, given by

$$-\sum p \ \log_2 p$$

Where p is the probability of an intensity in the normalized local neighborhood. In phase contrast images cells typically have a “rougher” texture than the background, leading to a higher entropy. The filtered images are Otsu thresholded and smoothed with a closing operation to define areas containing cells. Images adjacent in time are subtracted to create a difference mask– if a cell is present in one frame and detached in the next, it shows up as a continuous region in the mask that can be identified and registered as a detachment event. These continuous regions are thresholded by area to remove noise caused by small particles or optical effects.

The largest drawback to counting detachment events by difference is a relatively high false positive rate. There are two main mechanisms by which spurious detections of detachment events occur. First, small regions between cells may have a borderline entropy value. Whether they are thresholded or not seems to vary from frame to frame even if the positions of the cells around them do not change significantly. Second, cells may be dislodged from their resting position without being fully detached from the substrate. These false positives are addressed by user input. At the start of analysis for any experiment, the user is asked to outline the area of three small cells to set the area threshold for the difference mask. This is preferable to a fixed threshold because the typical cell size varies greatly between cell lines. The cell counting utility also has an optional mode where the user may verify any candidate detections of detachment events to manually reduce the false positive rate. It is generally recommended to employ manual user verification, although it is unnecessary for experiments with little debris in the field of view and well-separated cells.

**Agent-based Model of Differential Diffusion (Supplementary information S2)**

We developed an agent-based model to illustrate a postulated relationship between cell-matrix adhesion strength and migration speed. Each cell is represented by an agent with a cell-matrix adhesion strength positioned on a 2-D grid. At each time step, an agent randomly generates a propulsion force, sampled from a 2D Gaussian distribution. If the magnitude of this force is greater than the cell-matrix adhesion strength, the cell moves one grid space.

The following parameters were defined for each cell line:

σ – Standard deviation of the propulsion force distribution

φ – Cell-matrix adhesion strength

μ – Slope of mean squared displacement over time, i.e. 4D where D is the diffusion coefficient

The problem of tumor migration is considered over a space defined by a discrete grid, where each space is one cell diameter in size. Multiple cells may occupy the same coordinates on the grid. Cells are randomly initialised in a region of the grid specified by an initial tumor radius, the fractional area density of cells within that radius (may be greater than 1 to simulate crowding), and the initial proportion of each cell line. Each time step of the model represents 30 minutes of real time. During each time step, cells are selected in a random order and actions are performed. The cell moves randomly according to the following:

$$\left|\mathrm{F_p}\right| \sim \mathcal{R}(\sigma^2), \text{ where } \mathcal{R}(\sigma^2) \text{is the Rayleigh distribution}$$

$$\frac{F_p}{\left|F_p\right|} \sim \text{Random unit vector in Moore neighbourhood}$$

$$\text{Move one grid space in the in the direction } \frac{\boldsymbol{F_p}}{\left|\boldsymbol{F_p}\right|} \text{ if } \left|\boldsymbol{F_p}\right| > \phi.$$

Given values of μ and ϕ measured from our experiments, a value of σ can be chosen to match the simulation to the two values, making the model self-consistent. This comes from the fact that the adhesion strength effectively acts as a threshold on the success rate of motion. Using the cumulative distribution function of the Rayleigh distribution *P(F)*, a value of σ can be chosen as follows to ensure that the average distance travelled by cells over time *in silico* matches the experimentally observed value of μ:

$$P(F) = 1 - e^{-\frac{F^2}{2\sigma^2}}$$

Let the success rate of a jump in a given timestep $\Delta t$ be defined as $S$.

$$S = 1 - F(\phi)$$

The mean time required for a cell to move one square on the grid (i.e. one cell diameter $d$) is thus given by $\frac{\Delta t}{S}$ .

Hence for a given slope of mean squared displacement μ,

$$\frac{C\mu\Delta t}{S} = d^2$$

Where the constant $C$ accounts for the discretization of the movement directions and of the jump lengths in space. Substituting the equation for $S$ into the above relationship yields

$$\sigma^2 = -\frac{\phi^2}{\ln\frac{C\mu\Delta t}{d^2}}$$

*C* can be obtained from simulations by setting *S* to 1 (i.e. zero cell-matrix adhesion), allowing all cells to move one space per time interval. The expected slope of the mean squared displacement is then $\frac{d^2}{\mu}$, allowing *C* to be solved from the fitted value of μ.

In the simulations used in this work, $d$ was set to 19.83 μm to represent the average diameter of all cell lines studied. $\Delta t$ was set to 10 minutes, so that the success rate *S* for each cell was between 0 and 1 for values of $\sigma^2$ calculated using the experimental migration rates.

Supplementary Table 1: Differential diffusion model is self-consistent. Simulations (*n* = 100) were run using the analytically calculated $\sigma^2$ as described in the text, first assuming C = 1. μ
was determined by fitting the mean squared-displacement to a linear function of time in each line. The correct value of *C* was derived as described in the supplementary text. Then 100 simulations were run using $\sigma^2$ calculated with this value (*C* = 1.4924). After correction for *C*, we were able to recover values for μ close to the experimentally derived values in each cell line with a mean deviation of 5.5%.

| Line | Experimental μ | Simulated μ (C= 1.4924) |
|---|---|---|
| AS | 7.814 | 6.994 |

| | | |
|---|---|---|
| AN1 | 23.738 | 22.921 |
| AN2 | 20.951 | 20.861 |
| BS1 | 17.516 | 16.544 |
| BS2 | 14.364 | 13.548 |
| BW | 29.168 | 28.182 |
| CU1 | 24.760 | 23.900 |
| CU2 | 22.767 | 23.077 |
| CU3 | 24.610 | 28.182 |

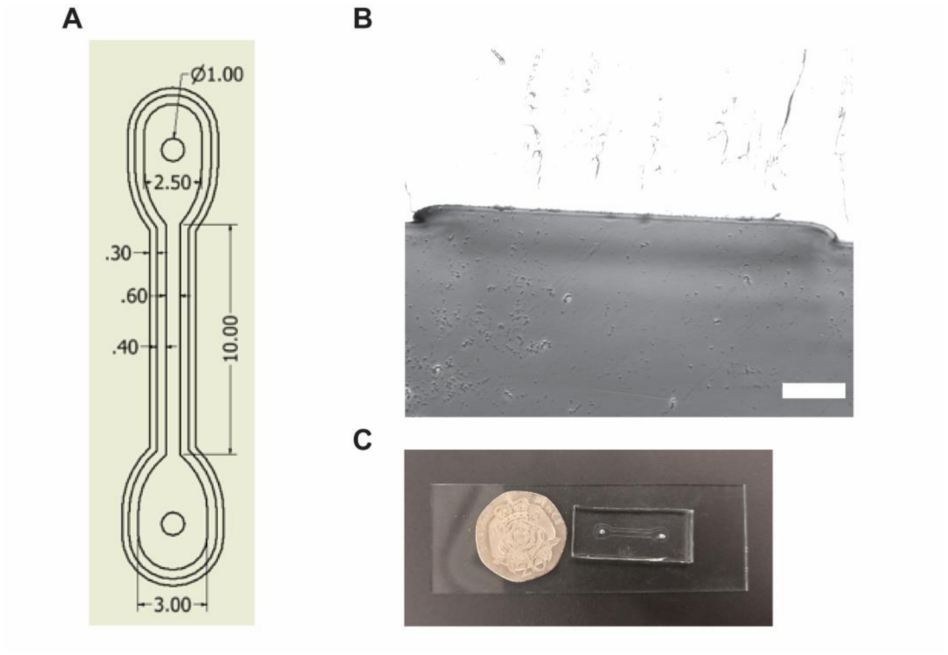


**Supplementary Figure 1: Microfluidic shear assay apparatus.**

(A) Channel dimensions.

(B) Channel cross-section. The average channel dimensions were (655.00 ± 21.26) µm × (28.00 ± 1.58) µm (W×H) (± sample SD).

(C) Device cast in PDMS and plasma-bonded to glass slide.

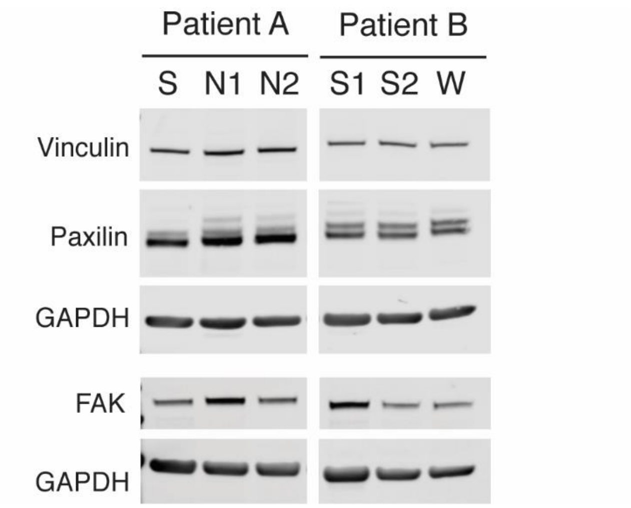

**Supplementary Figure 2: Western Blot performed on GBM primary cell lines**

Western blot assay was performed for all patients' cell lines. Representative western blot image showing the expression of vinculin, paxillin and FAK. GAPDH served as a loading control.

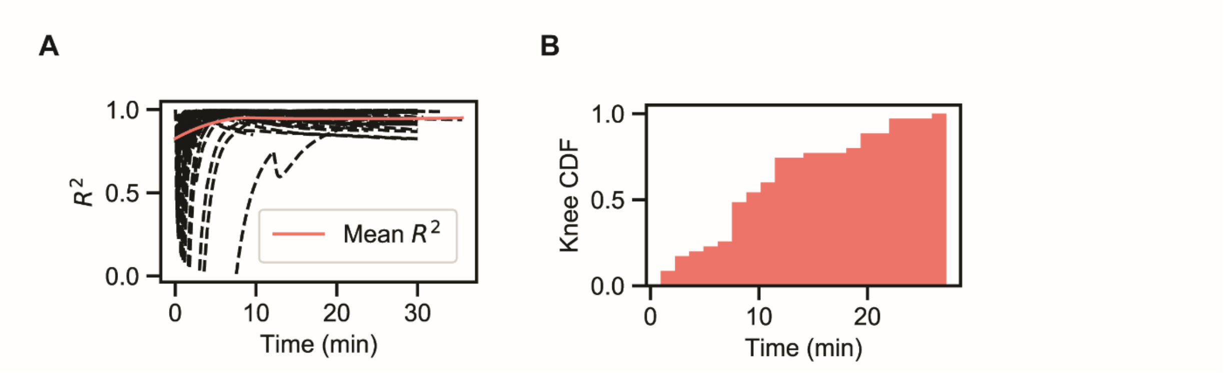


**Supplementary Figure 3: Determination of a cut-off for detachment observation.**

A knee in each curve was identified using two independent methods to find the time point at which detachment begins to slow down.

(A) Increasing windows of time were taken for a linear fit and the quality of the fit was assessed using $R^2$. The mean $R^2$ of all experiments reached a maximum at 8.74 minutes.

(B) The Kneedle algorithm was applied to each curve. 75% of experiments had a knee after 7.5 minutes according to this approach.

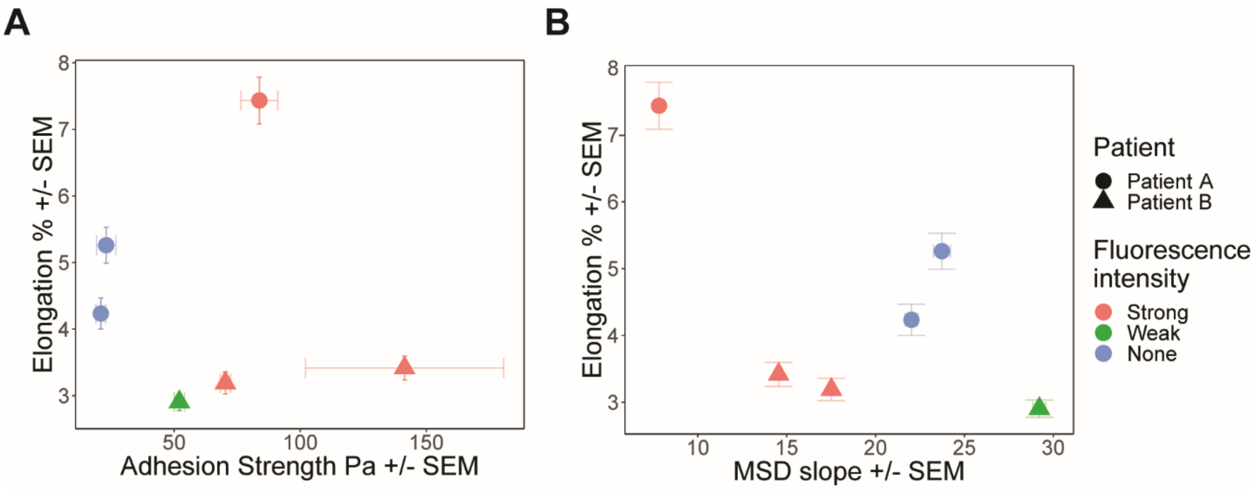


**Supplementary Figure S4: Relationship between elongation, adhesion and migration**

Cell migration and cell-matrix adhesion strength did not correlate with cell elongation.

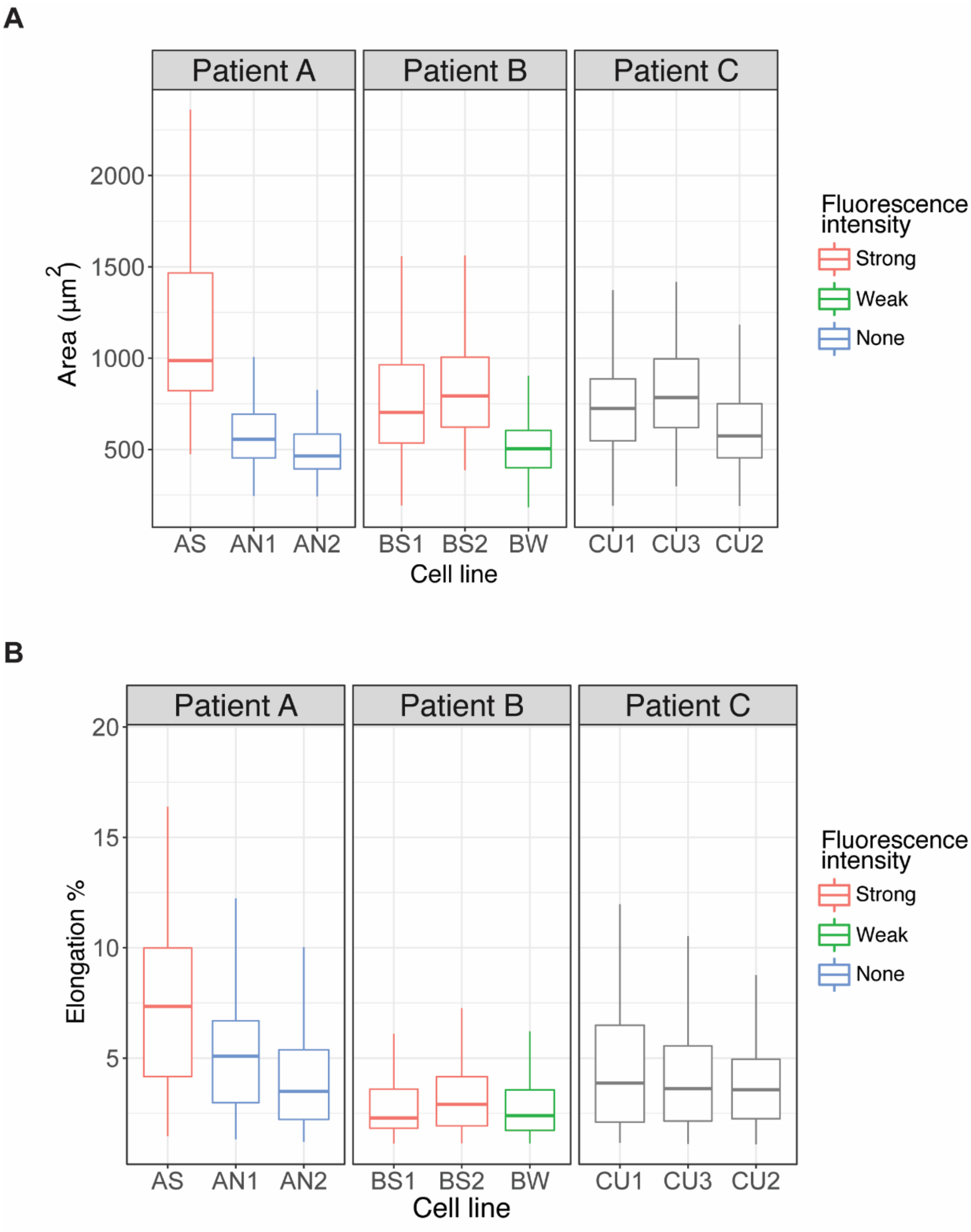


**Supplementary Figure 5: Morphological Characteristics of GBM Patients Derived Cell Lines**

Morphological quantification of cells cultured on PDMS substrates of ~1.5 kPa stiffness (n > 100 cells).

(A) Boxplots show spatial heterogeneity in cell area associated with sampling location and amount of 5-ALA fluorescence. Not accounting for sampling location and 5-Ala intensity show no significant difference in cell area between patient A, B and C. Significant difference was found between patients A, B (***$p<0.001$) and C (***$p<0.001$) when we accounted for sampling location and 5-ALA intensity. Patient C cells were derived from similar locations of unrecorded intensity within the patient's tumor.

(B) Boxplots of cell elongation in Patients A, B and C. Patient A exhibited a more elongated morphology compared to cells derived from patient B (***$p < 0.001$) and patient C (***$p < 0.001$).

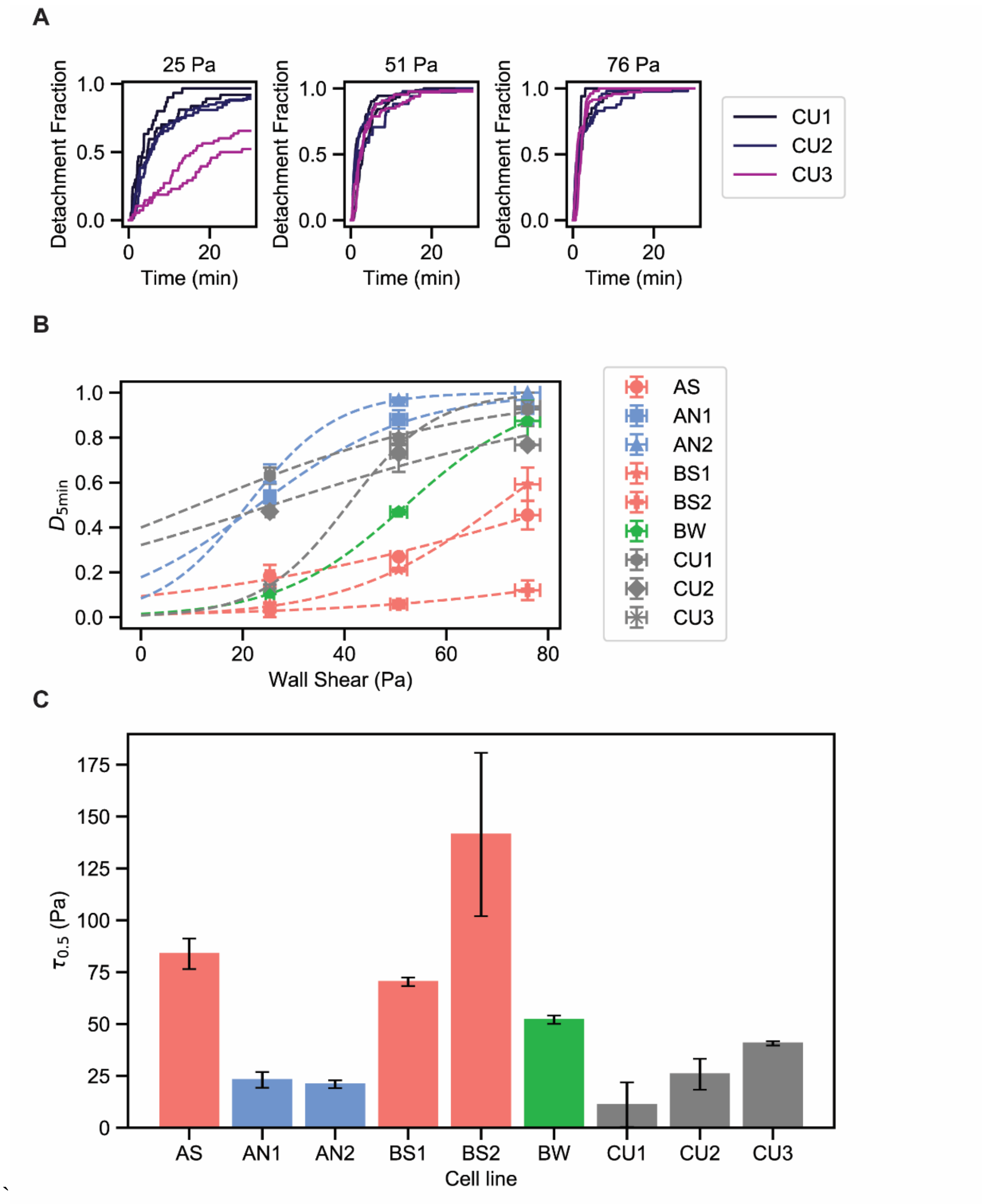


**Supplementary Figure 6: Characteristics of Patient C Cell-matrix Adhesion Strength**

(A) Time-detachment profiles for 3 cell lines derived from Patient C (n=2) for multiple shear stresses. Detachment increases with shear stress. Detachment initially occurs rapidly and then plateaus, consistent with Patients 1 and 2.

(B) The inflection point of each sigmoidal curve was extracted to define $\tau_d$, a measurement for the cell-matrix adhesion strength for all 9 cell lines derived from three Patients.

(C) Differences in cell matrix adhesion strength for all 9 cell lines. Cell lines from Patient C had adhesion strength similar to weak- or non-fluorescent lines.

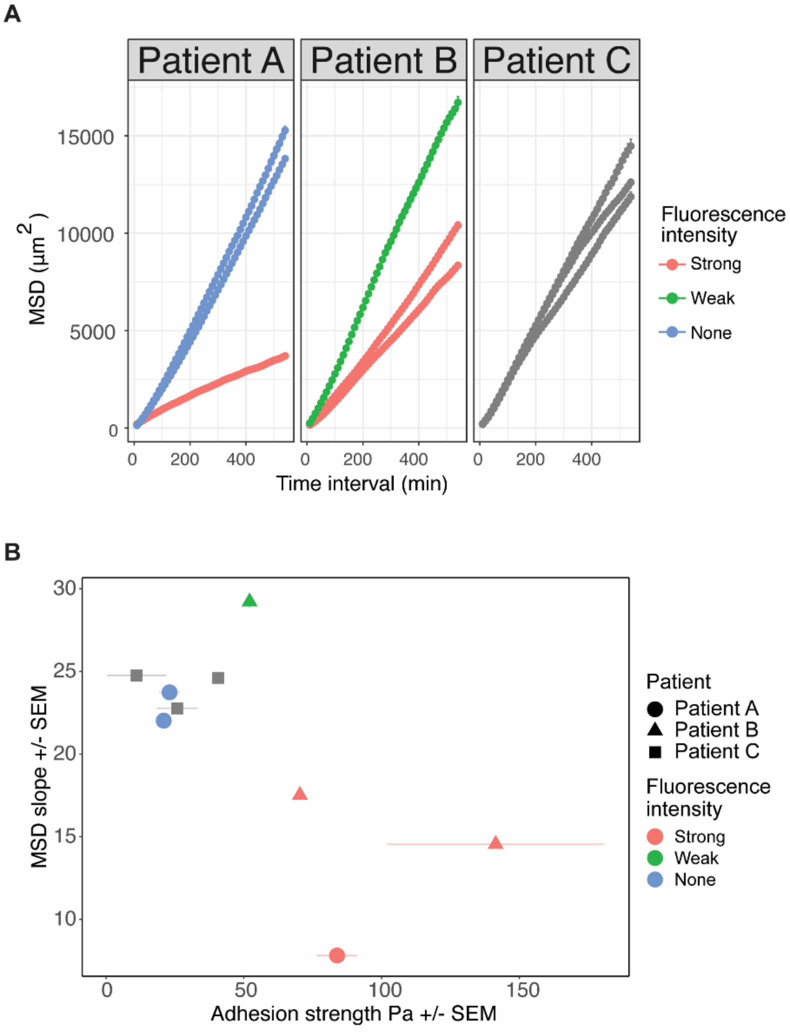


**Supplementary Figure 7: Relationship between Cell Migration and Cell-matrix Adhesion Strength**

A) Mean squared displacement of Nine cell lines derived from 3 patients calculated from cell trajectories with vertical bars representing standard errors.

B) More adhesive cells are less migratory as demonstrated by all patients' cell lines.

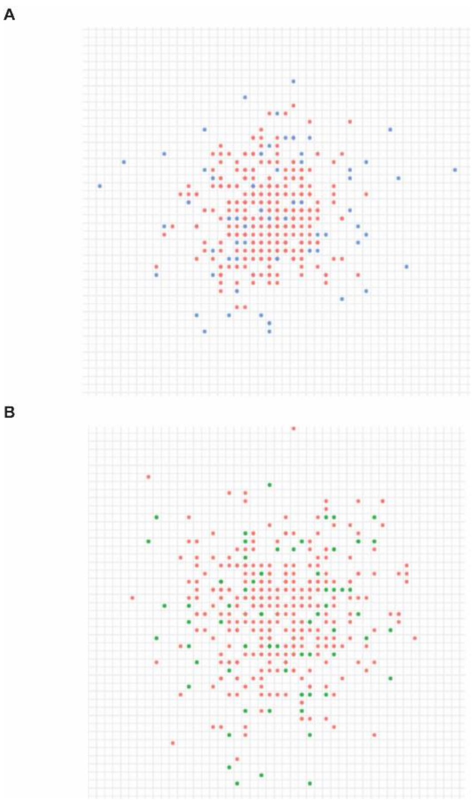


**Supplementary Figure 8: Agent based model simulating GBM mixed populations from core and marginal tumor cells**

Agent based model showing a mixture of weak- and strongly- fluorescent cell lines evolving over time. Left: AS (red) and AN1 (blue), Right: BS1 (red) and BW (green). The snapshots shown are simulations at 24 hours, with time interval 10 minutes and grid size 19.83 µm, assuming no cell division with an initial density of 5 cells per grid square and an initial tumor radius of 5 squares. Strongly-fluorescent cell lines are localized in the center of the tumor, and the weak-fluorescent lines in the margins.
